# Increasing molar activity by HPLC purification improves ^68^Ga-DOTA-NAPamide tumor accumulation in a B16/F1 melanoma xenograft model

**DOI:** 10.1101/537258

**Authors:** Jan Lennart von Hacht, Sarah Erdmann, Lars Niederstadt, Sonal Prasad, Asja Wagener, Samantha Exner, Nicola Beindorff, Winfried Brenner, Carsten Grötzinger

**Author notes:** Corresponding author: (CG). These authors contributed equally to this work (Jan Lennart von Hacht, Sarah Erdmann).

## Abstract

**Purpose:** Melanocortin receptor 1 is overexpressed in melanoma and may be a molecular target for imaging and peptide receptor radionuclide therapy. ^68^Gallium labeling of DOTA-conjugated peptides is an established procedure in the clinic for use in positron emission tomography imaging. Aim of this study was to compare a standard labeling protocol against the ^68^Ga-DOTA peptide purified from the excess of unlabeled peptide.

**Procedures:** The MC1R ligand DOTA-NAPamide was labeled with ^68^Ga using a standard clinical protocol. Radioactive peptide was separated from the excess of unlabeled DOTA-NAPamide by HPLC. Immediately after the incubation of peptide and ^68^Ga (95 °C, 15 min), the reaction was loaded on a C18 column and separated by a water/acetonitrile gradient, allowing fractionation in less than 20 minutes. Radiolabeled products were compared in biodistribution studies and PET imaging using nude mice bearing MC1R-expressing B16/F1 xenograft tumors.

**Results:** In biodistribution studies, the non-purified ^68^Ga-DOTA-NAPamide did not show significant uptake in the tumor at 1 h post injection (0.78% IA/g). By the additional HPLC step, the molar activity was raised around 10,000-fold by completely removing unlabeled peptide. Application of this rapid purification strategy led to a more than 8-fold increase in tumor uptake (7.0% IA/g). The addition of various amounts of unlabeled DOTA-NAPamide to the purified product led to a blocking effect and a decreased specific tumor uptake, similar to the result seen with non-purified radiopeptide. PET imaging was performed using the same tracers for biodistribution. Purified ^68^Ga-DOTA-NAPamide, in comparison, showed superior tumor uptake.

**Conclusions:** We demonstrated that chromatographic separation of radiolabeled from excess unlabeled peptide is technically feasible and beneficial, even for short-lived isotopes such as ^68^Ga. Unlabeled peptide molecules compete with receptor binding sites in the target tissue. Purification of the radiopeptide therefore improved tumor uptake.

## Introduction

Cutaneous malignant melanoma is one of the most lethal forms of cancer. Its incidence is increasing rapidly, making it a significant public health threat.[1] Melanocortin receptor 1 (MC1R) is overexpressed in most melanomas, making it a promising molecular target for diagnosis and peptide receptor radionuclide therapy (PRRT).[2, 3] Because of their low molecular weight, low immunogenicity and excellent tumor penetration, radiopeptides have attracted a steadily increasing interest in receptor-mediated tumor targeting.[4, 5]

Because of its enhanced spatial resolution and high sensitivity, positron emission tomography (PET) has been developed into a valuable diagnostic tool, particularly for the detection of small metastases. Since commercial ^68^Germanium/^68^Gallium generators became widely available, ^68^Ga labeling of chelator-conjugated peptides turned into an established clinical procedure for use in PET imaging.[6] Due to its short half-life (67.71 min), ^68^Ga has a higher molar activity (lower mass/activity ratio) than other nuclides in nuclear medicine, resulting in an unfavorable reaction stoichiometry.[7] In the radiochemical chelation process of ^68^Ga incorporation into 1,4,7,10-tetraazacyclododecane-1,4,7,10-tetraacetic acid (DOTA), a high molar excess of DOTA-conjugated peptide over radiometal is usually used to attain high ^68^Ga complexation yields. The applied excess of cold peptide mass for ^68^Ga chelator loading typically ranges from 1,000-fold to 10,000-fold. This fraction is usually not removed before injection into the patient and it will compete for binding sites at the tumor, resulting in lower detection sensitivity. The aim of this study was to compare a standard labeling protocol against a labeling and HPLC-purification protocol, which removes the excess of unlabeled peptide. We investigated the influence of peptide mass/molar activity on tumor accumulation of the MC1R ligand ^68^Ga-DOTA-NAPamide.

## Materials and Methods

### Peptides

DOTA-NAPamide was from ABX (Radeberg, Germany), [Nle^4^,d-Phe^7^]-melanocyte-stimulating hormone (NDP-MSH) from peptides&elephants (Hennigsdorf, Germany). Peptides were analyzed by a Finnigan Surveyor MSQ Plus LC-MS (Thermo Finnigan, Bremen, Germany) to confirm the presence of the correct molecular mass. Peptides were used at a purity of greater than 95%.

### Competitive Binding and Saturation Assays

[Nle^4^,d-Phe^7^]-melanocyte-stimulating hormone (NDP-MSH) was iodinated with Na^125^I by the chloramine-T method and it was purified from unlabeled peptide by HPLC as described below. Saturation and competitive binding studies were performed with live cells. 40.000 B16/F1 cells were seeded per well into a 96-well flat bottom cell culture plate. For competitive binding, medium was removed and 50 μL of binding buffer (50 mM 4-(2-hydroxyethyl)-1-piperazineethanesulfonic acid (HEPES) pH 7.4, 5 mM MgCl_2_, 1 mM CaCl_2_, 0.5% BSA, protease inhibitor cocktail cOmplete [Roche Applied Science, Penzberg, Germany]) with increasing concentrations of non-radioactive peptide was added to the cells. Additionally, 50 μL binding buffer with 100,000 cpm of ^125^I-[Nle^4^,d-Phe^7^]-MSH was added. After 30 minutes of incubation at 37 °C, cells were washed 4 times with cold washing buffer (50 mM Tris-HCl pH 7.4, 125 mM NaCl, 0.05% BSA). 100 μL 1 N NaOH was added added to lyse cells. Lysates were transferred into vials and measured using a gamma counter (Wallac 1470 Wizard, PerkinElmer, Waltham, MA, USA). The saturation assay was performed by adding 100 μL of binding buffer with increasing amounts of ^125^I-NDP-MSH to the cells in the presence or absence of 1 μM of unlabeled NDP-MSH.

### Radiolabeling of DOTA-NAPamide

Radiolabeling experiments were performed on a Modular Lab PharmTracer synthesis module (Eckert & Ziegler, Berlin, Germany) which allows fully automated cassette-based labeling of gallium tracers utilizing a pharmaceutical grade ^68^Ge/^68^Ga generator (GalliaPharm, 1.85 GBq, good manufacturing practice (GMP)-certified; Eckert & Ziegler GmbH, Berlin, Germany). Cassettes were GMP-certified and sterile. They were used without pre-conditioning of the cartridges. Gallium generator at three months post calibration was eluted with aqueous HCl (0.1 M, 7 ml) and the eluate was purified on an ion-exchange cartridge followed by elution using 1 m of 0.1 M HCl in acetone. An aliquot of DOTA-NAPamide, 50 μg (stock solution 1 mg/ml in 10% DMSO, 90% water) was mixed with 500 μl 0.1 M HEPES buffer (pH 7) and heated for 500 s at 95 °C. After the reaction, the reactor was cooled with 500 ml of saline and without post-processing. The contents of the reactor were directly used for subsequent HPLC purification.

### HPLC Purification

^68^Ga-labeled peptide was separated from non-radioactive NAPamide by reverse phase HPLC on an Agilent 1200 system (Agilent, Waldbronn, Germany). The complete mixture from the reaction chamber was loaded on an Eclipse XDB-C18 bonded silica column (Agilent, Waldbronn, Germany) and eluted with a linear gradient of acetonitrile (gradient 15 – 45% B in A over 20 min, flow rate of 1 ml/min, solvent A water + 0.1% trifluoroacetic acid (TFA), solvent B acetonitrile + 0.1% TFA, column at 55 °C). ^68^Ga-DOTA-NAPamide was detected using a FlowStar LB513 detector (Berthold, Bad Wildbad, Germany) equipped with a BGO-X (5 μl) chamber. The unlabeled peptide moiety was detected via a diode array absorbance detector. ^68^Ga-DOTA-NAPamide was separated with the help of an automated fraction collector.

### *In vivo* Biodistribution Assays

B16/F1 cells (3×10^6^) were inoculated subcutaneously on the right shoulder of NMRI-*Foxn1*^*nu*^ /*Foxn1*^*nu*^ mice (Janvier Labs, Saint-Berthevin, France). After 1-2 weeks, tumor bearing mice were injected with approximately 5 MBq of Ga^68^-DOTA-NAPamide to the tail vein via a catheter. Mice were sacrificed and dissected 1 h after injection. The B16F1 tumor, blood, stomach, pancreas, small intestine, colon, liver, spleen, kidney, heart, lung, muscle and femur samples were weighed and uptake of radioactivity was measured by a gamma counter (Wallac 1470 Wizard, Perkin Elmer, Waltham, MA, USA). To determine the effect of unlabeled ligand on the tumor uptake, either 0.5 nmol or 0.05 nmol non-labeled DOTA-NAPamide was co-injected.

### *In vivo* PET/MRI Imaging

The study protocol was approved by the local committee for animal care according to the German law for the protection of animals. All applicable institutional and national guidelines for the care and use of animals were followed. Positron emission tomography (PET) / magnetic resonance imaging (MRI) (1 Tesla nanoScan PET/MRI Mediso, Hungary) was performed at the Berlin Experimental Radionuclide Imaging Center (BERIC), Charité – Universitätsmedizin Berlin. Anatomic MRI scans were acquired using a T2-weighted 2D fast spin echo sequence (T2 FSE 2D) with the following parameters: coronal sequentially, matrix 256×256×20 with dimensions 0.36×0.36×1.5mm^3^, TR: 8695 ms, TE: 103 ms, and a flip angle of 180°. PET scans were performed for 90 min starting directly before intravenous injection of 0.15 mL of tracer, corresponding to a ^68^Ga activity of approximately 15 MBq).). PET images were reconstructed from the raw data with the following image sequence: 6 × 10 s, 6 × 30 s, 5 × 60 s and 8 × 600 s. The tracer standardized uptake value (SUV) in the tumor tissue was determined by manual contouring of a volume of interest (VOI) of the PET images using PMOD 3.610 (PMOD Technologies, Zürich, Switzerland).

## Results

### DOTA-NAPamide Binds to MC1R in vitro

To confirm MC1R expression in the melanoma cell line B16/F1 and to assess the affinity of DOTA-NAPamide towards the receptor, competitive radioligand binding and saturation binding assays using ^125^I-labeled NDP-MSH were performed (Fig 1A). DOTA-NAPamide showed a high affinity for the murine MC1R expressed in the B16/F1 cells, with a calculated Ki of 0.37 nM from and a K_D_ of 660 pM. To exclude a negative effect of gallium (Ga) incorporation into the DOTA chelator on binding affinity, competitive in vitro binding assays were performed using either DOTA-NAPamide or DOTA-NAPamide complexed with non-radioactive Ga. Figure 1B shows nearly identical concentration-response curves and K_i_ values (0.40 nM vs. 0.43 nM) for unlabeled and Ga-labeled DOTA-NAPamide binding to B16/F1 cells. This demonstrated that chelation with Ga did not affect the affinity of DOTA-NAPamide towards the MC1R receptor.

**Fig 1:**
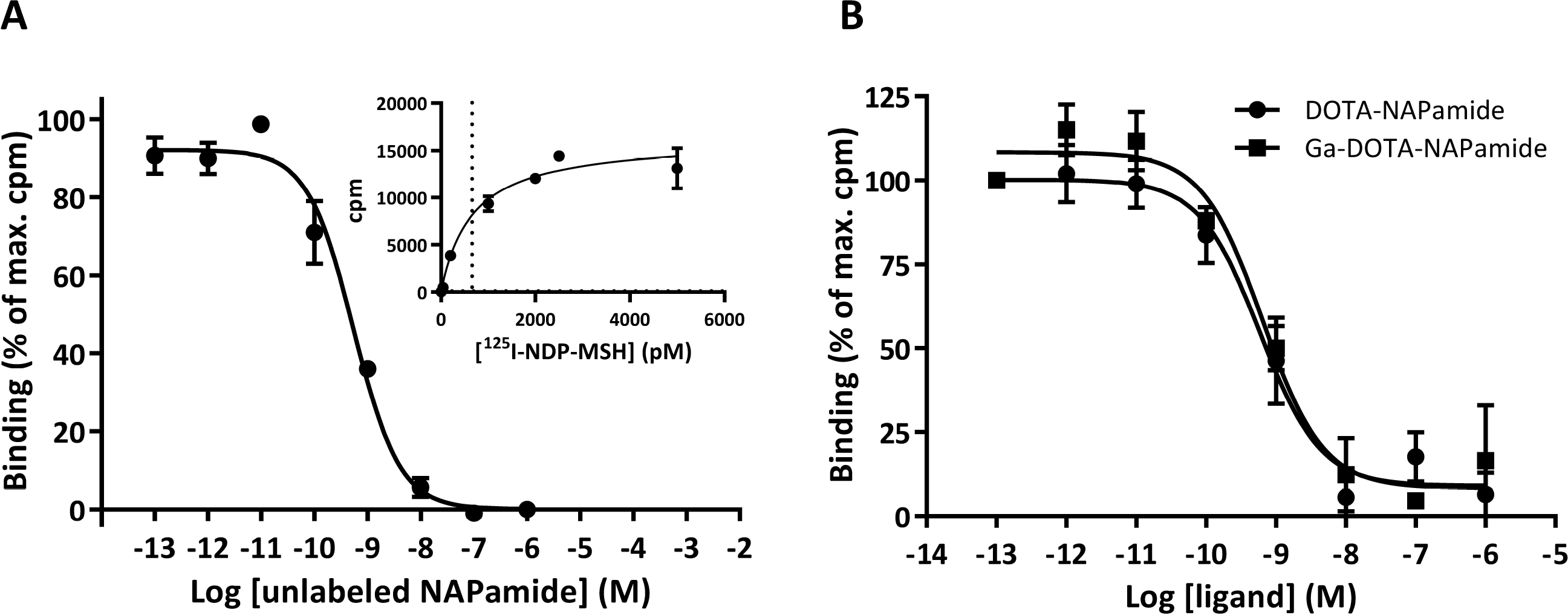
DOTA-NAPamide and Ga-DOTA-NAPamide are high affinity ligands for the melanocortin 1 receptor (MC1R): ^125^I-NDP-MSH ligand displacement and saturation assays performed with whole B16/F1 cells. **A** Various concentrations of NAPamide were used to displace ^125^I-NDP-MSH. The inset shows a saturation experiment to determine the dissociation constant. **B** ^125^I-NDP-MSH ligand displacement assay to assess the impact of gallium chelation on DOTA-NAPamide. Curve fits were performed in GraphPad Prism by applying a one‐site binding equation. (n=3; mean +/− SEM)

### Unlabeled DOTA Peptide Can Be Removed by HPLC

For removal of unlabeled excess of DOTA-NAPamide and of non-incorporated ^68^Ga after completion of the radiochemical reaction, the product was transferred from the reactor to an HPLC equipped with a C18 reverse-phase column and a fraction collector. Figure 2A shows an exemplary separation run for ^68^Ga-DOTA-NAPamide and unlabeled DOTA-NAPamide. Free ^68^Ga does not exhibit a distinct interaction with the C18 column and elutes close to the dead time of the HPLC. For the labeling reaction, the peptide was used in an excess compared to ^68^Ga (10,000:1 molar ratio) and it showed a slightly prolonged retention time in comparison to ^68^Ga-DOTA-NAPamide. This was exploited to purify ^68^Ga-DOTA-NAPamide (Fig 2A). To demonstrate that an excess of unlabeled peptide would displace ^68^Ga^68^-DOTA-NAPamide from its receptor, the radiotracer was incubated on B16/F1 cells in vitro either alone or with a 1,000-fold or 10,000-fold excess of DOTA-NAPamide (Fig 2B). Indeed, both concentrations of unlabeled peptide were able to displace the radiopeptide as compared to incubation with buffer. In comparison to purified tracer alone, a 1,000-fold excess led to an approximately 20% decrease and a 10,000-fold excess of unlabeled peptide diminished the overall binding to less than 50% (Fig 2B).

**Fig 2:**
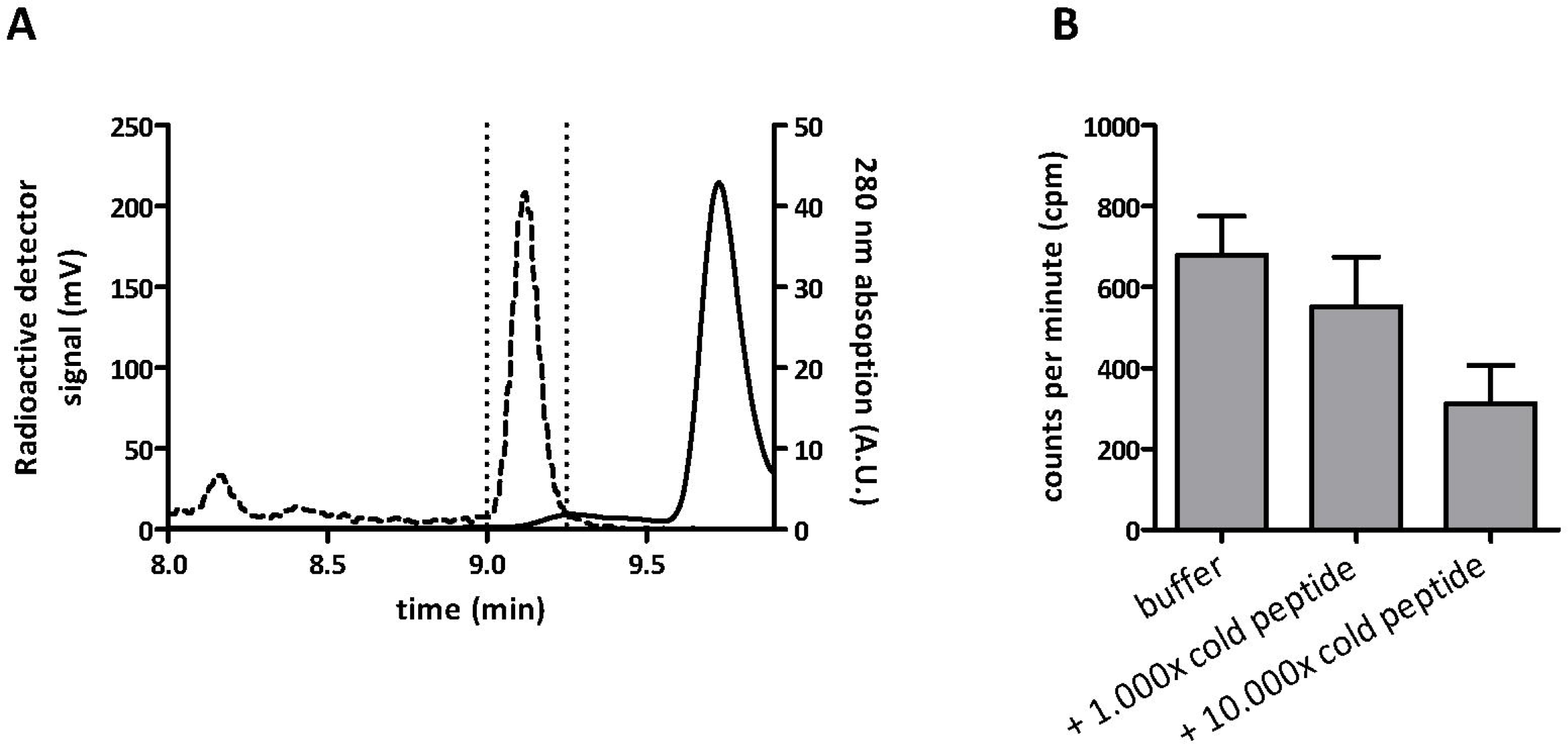
**A** Reverse-phase HPLC chromatogram of a ^68^Ga-DOTA-NAPamide purification. The dashed line shows the radiodetector signal and the peak there represents the ^68^Ga-DOTA-NAPamide fraction, while the solid line shows the 280 nm absorption signal with the peak of DOTA-NAPamide. The area surrounded by dotted lines represents the purified fraction used in in-vivo experiments. **B** In-vitro displacement of ^68^Ga-DOTA-NAPamide from B16/F1 cells by an excess of unlabeled DOTA-NAPamide (n=3; mean +/− SEM).

### Purification of the Tracer Leads to an Improved Tumor Uptake

Fig 3 shows the results of an in-vivo biodistribution experiment in B16/F1 xenograft-bearing mice 1 hour after i.v. injection of ^68^Ga-DOTA-NAPamide. The non-purified tracer from the standard procedure showed a very low uptake into subcutaneously grown B16/F1 tumor xenografts (0.78% IA/g). Removal of unlabeled DOTA-NAPamide led to a more than 8-fold increase in tumor uptake, with 7.0% IA/g in the tumor for the purified ^68^Ga-DOTA-NAPamide. Except for a moderate uptake in the kidneys, all other tissues showed only a small uptake of the tracer, mostly below 0.5% IA/g. In animals treated with ^68^Ga-DOTA-NAPamide produced by the standard procedure, the kidney uptake was slightly higher compared to the purified tracer (4.56% IA/g vs. 3.07% IA/g) (Fig 3).

**Fig 3:**
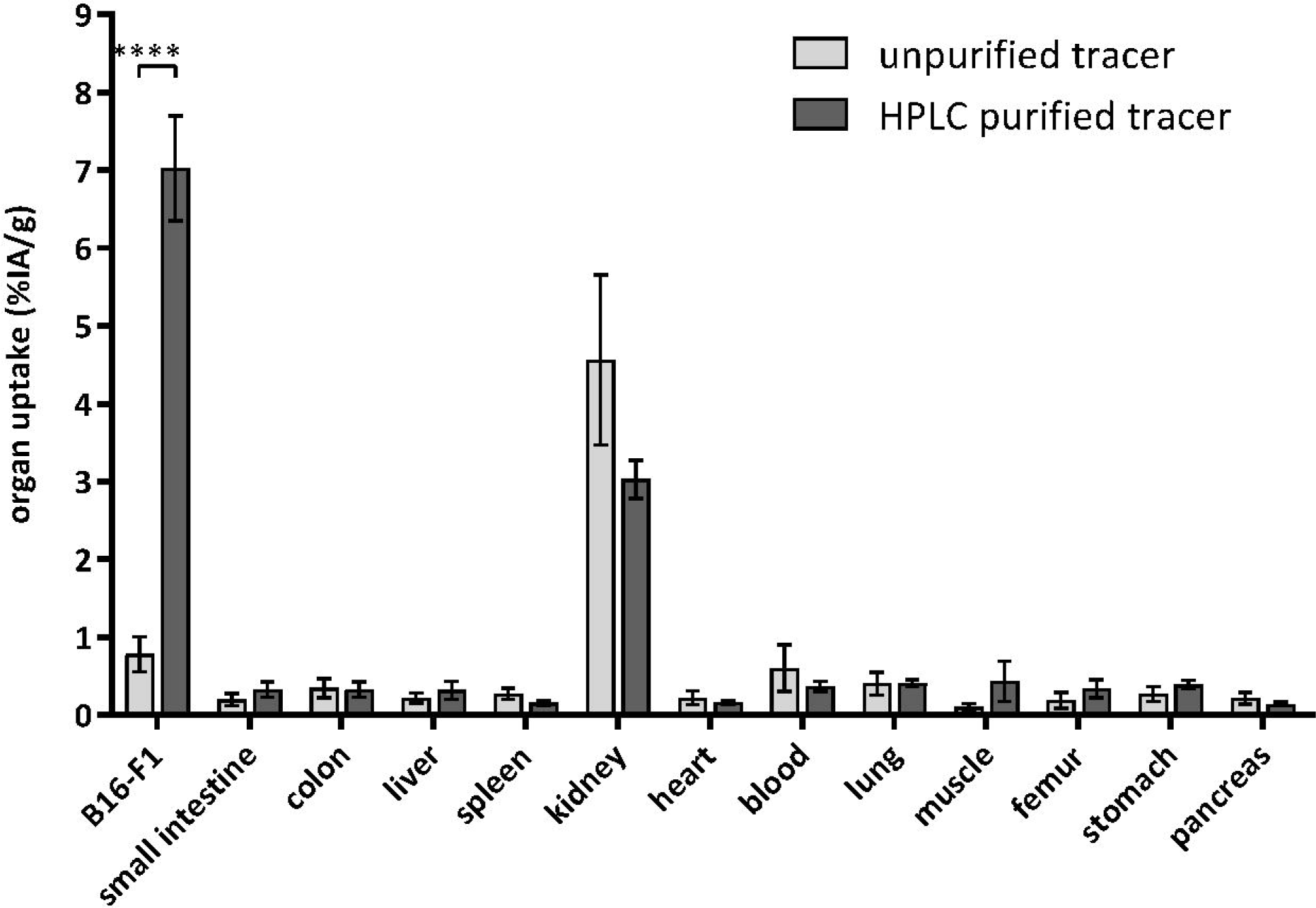
Biodistribution at 1 h after i.v. injection of approximately 5 MBq radiotracer along with approximately 0.5 μg peptide. Results of ^68^Ga-DOTA-NAPamide produced by the standard protocol (n=7) are shown in light gray, and results obtained with the purification protocol (n=7) are shown in dark gray. (mean +/ SEM; ****p= 0.0001)

### Tumor Uptake Enhancement is Reversed by Coinjection of Unlabeled Peptide

The effect of cold peptide mass was studied by injecting a constant amount of purified ^68^Ga-DOTA-NAPamide together with different amounts of unlabeled DOTA-NAPamide. Mice were injected with 5 MBq (~50 fmol) alone or with an additional 1,000-fold (50 pmol) or 10,000-fold (500 pmol) excess of unlabeled peptide. This 10,000-fold excess of unlabeled peptide corresponds to the molar ratio in the standard protocol. The coinjection of DOTA-NAPamide led to significant loss of uptake in the MC1R-expressing B16/F1 tumor xenografts (Fig 4). While purified ^68^Ga-DOTA-NAPamide showed an uptake of 6.7% IA/g, coinjection of 50 pmol cold DOTA-NAPamide decreased the total uptake to 3.8% IA/g. 500 pmol coinjected peptide led to an approximately 6-fold decrease to 1.1% IA/g (Fig 4).

**Fig 4:**
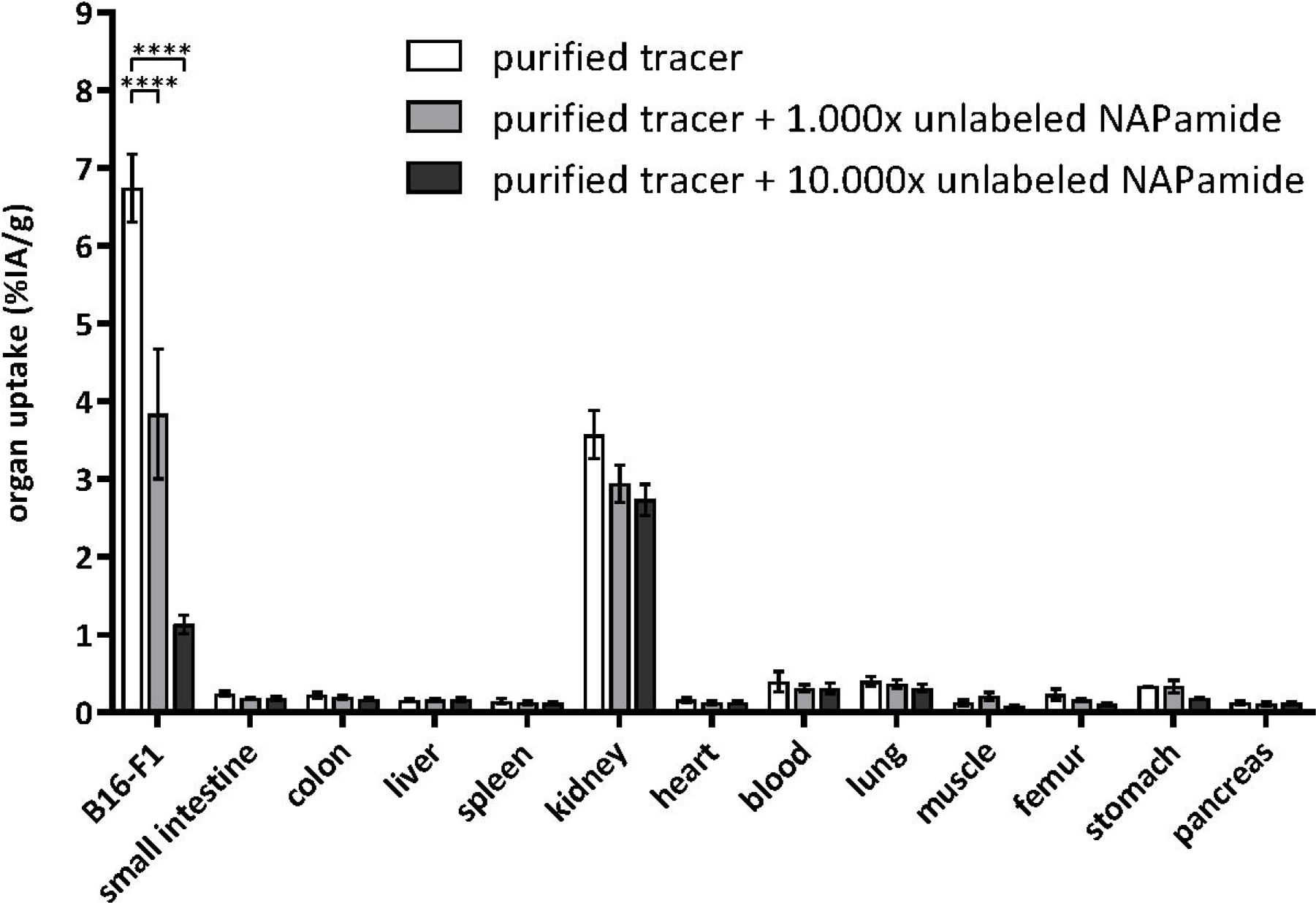
^68^Ga-DOTA-NAPamide biodistribution at 1 h after i.v. injection of approximately 5 MBq radiotracer. Purified ^68^Ga-DOTA-NAPamide was injected either alone or in parallel with an excess of unlabeled DOTA-NAPamide (1,000-fold or 10,000-fold excess over radiotracer). (n=3 per group; mean +/− SEM; ****p= 0.0001)

### PET imaging confirms the results of biodistribution studies

Mice bearing subcutaneous B16/F1 tumors on their right shoulder were injected with approximately 15 MBq ^68^Ga-DOTA-NAPamide in different formulations. Each mouse was given a different DOTA-NAPamide composition (A standard procedure, B purified, C purified + 1,000-fold excess of cold peptide, D purified + 10,000-fold excess of cold peptide). Dynamic PET images were taken from 5 to 90 minutes after injection. Fig 5 shows all four tested conditions at 1 h post injection. The tumor showed an increase in uptake of ^68^Ga-DOTA-NAPamide for the purified and the purified + 1,000-fold excess of cold peptide conditions (Fig 5 B and C). For the injections with a 10,000-fold excess of unlabeled peptide, the tracer uptake was on a similarly low level as for the unpurified tracer (Fig 5 A and D). Since the peptide is excreted through the kidneys into the bladder, a high signal was observed in both organs (kidneys not visible in Fig 5 due to their location in a different plane).

**Fig 5:**
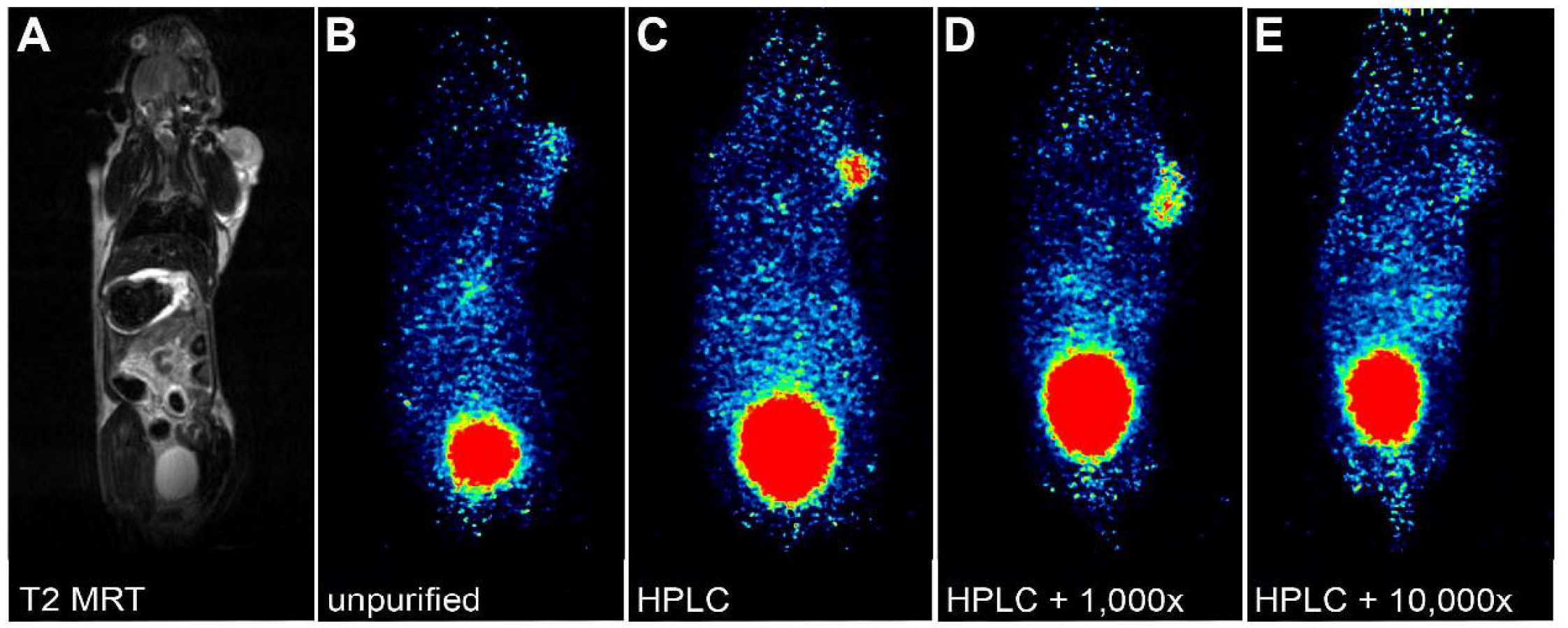
Coronal sections from MRI and PET scans of mice bearing B16/F1 tumors on the right shoulder at 1 h after tail vein injection of ^68^Ga-NAPamide with various ratios of labeled and unlabeled peptide. **A** T2-weighted MRI (corresponding to PET image in B) **B** unpurified ^68^Ga-DOTA-NAPamide **C** HPLC-purified ^68^Ga-NAPamide **D** HPLC-purified ^68^Ga-NAPamide + 1,000-fold excess of DOTA-NAPamide, and **E** HPLC-purified ^68^Ga-NAPamide + 10,000-fold excess of DOTA-NAPamide.

### Tracer kinetics

To obtain kinetic profiles, mice were imaged for up to 90 minutes after injection of either of the four different tracer formulations. Images were reconstructed over short intervals: 10/30/60 seconds until 10 minutes p.i. and over 10 minutes thereafter. Fig 6A shows the kinetics for the purified ^68^Ga-DOTA-NAPamide. After defining volumes of interest (VOIs) for the tumor in each mouse, the resulting standardized uptake values were plotted over time from dynamic PET data. Early tracer kinetics in the tumor confirm the advantage of tracer purification: in the first 5 - 10 minutes after injection, the tracer reaches a maximum level in the tumors for all tested conditions (Fig 6B). Thereafter, uptake was slowly decreasing over the next 90 minutes. In the animal treated with a 1,000-fold excess of cold peptide mass, the initial amount of radioactivity in the tumor is slightly higher (SUV 0.63) than in the other tumors (mean SUV 0.43). Of note, the slope of tracer decrease in B16/F1 tumors was less steep for the purified ^68^Ga-DOTA-NAPamide.

**Fig 6:**
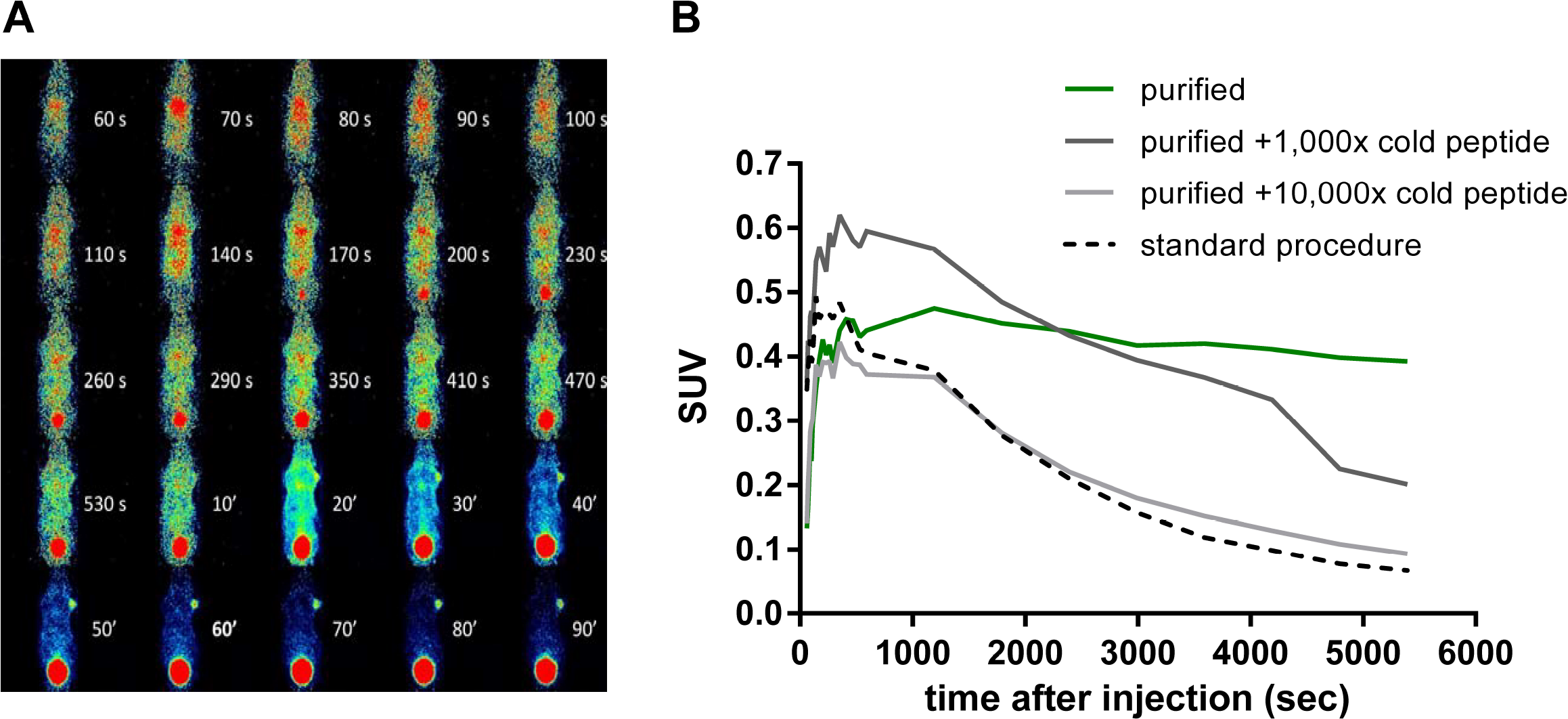
**A** Short-interval reconstruction from dynamic PET data of a mouse bearing a B16/F1 tumor on the right shoulder after tail vein injection of 5 MBq purified ^68^Ga-NAPamide. Injected peptide amounts: 0.34 nmol (unpurified and 10,000-fold excess), 3.4 nmol (1,000-fold excess) and 3.4 pmol (purified). **B** Time-activity relationships of ^68^Ga-DOTA-NAPamide B16/F1 tumor accumulation derived from dynamic PET data obtained after injection of four different tracer formulations (n=1).

## Discussion

The feasibility of targeting metastatic melanoma with MC1R ligands for nuclear medicine has been studied as early as 1990 and has since resulted in a multitude of ligand-chelator conjugates applied for biodistribution, SPECT and PET experiments.[8, 9] Froidevaux et al. developed the 8mer metabolically stable high-affinity MC1R ligand DOTA-NAPamide that showed favorable biodistribution and tumor uptake values ranging from 7.56% to 9.43% IA/g at 4 h p.i.[10] The radionuclides used were of moderate half-life (2.81 d for ^111^In, 3.26 for ^67^Ga). Another study with ^64^Cu-DOTA-NAPamide reported ~4% IA/g at 4 h p.i.[11] Only recently, one study used the short-lived ^68^Ga with this tracer demonstrating tumor demarcation in PET imaging yet not reporting biodistribution data for the B16/F10 tumors used.[12] However, no clinical study has been reported for the use of MC1R analogs in melanoma imaging so far.

In experiments preceding this study, we performed biodistribution studies using the MC1R ligands ^68^Ga-DOTA-NDP-MSH and ^68^Ga-DOTA-NAPamide in the B16/F1 melanoma model, yet failed to achieve the high tumor uptake previously reported for the longer-lived ^111^In and ^67^Ga complexes. We hypothesized that the unfavorable stoichiometry of the chelation reaction of the peptide conjugate with ^68^Ga and the resulting high excess of unlabeled ligand prevented higher tumor uptake due to competition for MC1R binding sites. Therefore, in the current study we aimed to compare a standard ^68^Ga labeling protocol against a labeling and HPLC-purification protocol, which removes the excess of unlabeled peptide. We investigated the influence of peptide mass/molar activity on tumor accumulation of the MC1R ligand ^68^Ga-DOTA-NAPamide using biodistribution and PET imaging.

For the standard labeling protocol used in this study, the product of 350 MBq ^68^Ga-DOTA-NAPamide (3.4 pmol ^68^Ga) is accompanied by 34 nmol (50 μg) of unlabeled peptide, resulting in a 10,000-fold excess over the radioligand. Due to this stoichiometry, the theoretical molar activity of 103.000 GBq/μmol for pure ^68^Ga-DOTA-NAPamide is diminished to an effective molar activity of just 10 Gbq/μmol. Lowering the peptide amount in the chelation reaction was no remedy as it dramatically reduces the radiochemical yield.

HPLC purification of the radiopeptide, however, improved tumor uptake in biodistribution studies by a factor of more than 8-fold, resulting in a tumor-to-kidney ratio of 2.33 and tumor-to-tissue ratios of better than 15 for all organs investigated (Fig 3). While others have argued HPLC purification was not convenient in clinical practice and developed solid phase extraction of ^68^Ga-exendin as an alternative method [13], we found using an HPLC to fractionate tracer was feasible and efficient. As most radiochemical laboratories have this instrumentation in place for routine quality control, cost may not be limiting. The procedure can also be performed rapidly - within 15 to 20 min. Similar successful approaches to purify ^11^C, ^18^F and ^67^Ga tracers have been reported.[14–16] However, with production of ^68^Ga tracers shifting towards kit preparation, HPLC purification may not have much clinical future. In addition, the enhancement of molar activity may not be of critical importance for the imaging of humans as the applied amount of activity (and thus, peptide) is several orders of magnitude lower than for rodents.

The appropriate amount of cold peptide mass or the best molar activity for optimal tumor imaging has been a matter of investigation for more than two decades now. Using the somatostatin receptor ligand ^111^In-pentetreotide, an early study by Breeman et al. found moderate molar activities (0.6, 6.0 MBq/μg) to yield the highest uptake in rat pancreas, while higher and lower values showed reduced uptake.[17] However, the majority of subsequent studies demonstrated that for efficient receptor targeting, low peptide doses should be administered. De Jong et al e.g. systematically investigated the effects of varying peptide mass at constant or varying molar activity on organ activity using ^111^In-[DOTA^0^,Tyr^3^]octreotide in tumor-bearing rats.[18] This study identified a principal detrimental effect of increasing peptide doses in both experimental settings with pancreas and tumor only showing limited benefit of an increased peptide dose of 0.2 (pancreas) or 0.5 μg (CA20948 tumors). Increasing amounts of radiolabeled peptide at constant molar activity were also used investigating the ^67^Ga-labeled GRPR ligand BZH3.[19] A peptide amount of 15 pmol per mouse showed highest tumor uptake (10.9% IA/g), with 5, 45 and 135 pmol yielding somewhat lower results (7.98, 8.0, 6.25% IA, respectively). Of note, intestines showed a continuously decreasing % IA/g uptake with increasing amounts of peptide, suggested to be due to increasing saturation of receptors. With limited molar activity of the ^68^Ga tracer, significant peptide dose reduction was only feasible at the expense of very small activities, insufficient for high-quality imaging in mice.[19]

Brom et al. compared the effect of co-injected unlabeled peptide over 5 orders of magnitude in in vivo biodistribution studies.[13] Low doses of unlabeled exendin-3, in the molar range of co-injected radioactive [Lys^40^(^111^In)]exendin-3 did not show a clear effect on tumor uptake. Eventually, the addition of 0.3 μg (~60 pmol) cold peptide led to a decrease in tumor uptake of the radioactive tracer. This dose was approximately 300-fold higher than the administered amount of ^111^Indium-labeled exendin. The same group compared the effect of co-injected unlabeled DOTA-minigastrin over four orders of magnitude (0.1 – 100 μg) in in vivo biodistribution studies. They observed an increasing drop in tumor uptake of their labeled DOTA-minigastrin for all tested amounts of cold co-injections (50 pmol - 50 nmol).[20] Using the standard labeling procedure in the current study, 500 pmol of unlabeled peptide were injected with an activity of 5 MBq ^68^Ga-DOTA-NAPamide. A detrimental effect of higher peptide dose had also been described in the first study describing DOTA-NAPamide as an MC1R tracer: tumor uptake at 170 pmol or 420 pmol peptide was much lower than with just 20 pmol.[10]

Chelators with higher labeling efficiency than DOTA may represent an alternative approach to obtaining high molar activities. Recently, the effect of varying molar activities (0.5 - 1,000 GBq/μmol) was studied for the two integrin-targeting ^68^Ga-TRAP peptides ^68^Ga-aquibeprin and ^68^Ga-avebetrin in biodistribution and PET imaging.[12] With this chelator (TRAP), very high molar activities (up to 5,000 GBq/μmol) could be produced during radiochemical labeling without additional HPLC purification.[21] Again, highest tumor uptake was found with highest molar activities with similar tumor-to-kidney ratios for all tested conditions. However, tumor-to-muscle ratios reached a maximum at 6 nmol (~3.5 GBq/μmol), indicating a benefit for moderate molar activities when muscle signal is limiting.[12] A variety of factors may lead to this, including target expression levels in muscle and other organs. Further studies will have to elucidate the relationship between expression, biodistribution and optimal molar activity for tumor imaging.

## Conclusions

Choosing the right molar activity and finding a way to obtain it in an efficient way remains to be a challenge during the introduction of new tracers. We showed that separation of radiolabeled and cold peptide via HPLC is technically feasible and beneficial, even for short-lived isotopes such as ^68^Ga. Unlabeled peptide molecules can strongly compete with receptor binding sites in the target tissue. Purification of the radiopeptide improved tumor uptake in in biodistribution studies and PET/MRI scans. Production of higher molar activity radiotracers may not only be important for imaging, but may also improve uptake and thus efficacy in therapeutic procedures such as PRRT.

## Acknowledgements

This work was supported by a grant from the German Ministry of Education and Research (BMBF grant IPT614A to CG), and in part by the Deutsche Forschungsgemeinschaft (DFG) for PET/MRI use (INST 335/454-1FUGG).

